# Bioinformatics analysis for L polymerase homology of arenaviruses

**DOI:** 10.1101/178012

**Authors:** Hasan Alrashedi, Ian Jones, Geraldine Mulley, Benjamin W Neuman

## Abstract

Many pathogenic viruses can transmit between humans and animals as zoonotic infectious often to cause dangerous disease with obvious clinical signs, sometimes globally. However, these outbreaks are generally only dealt with seriously when the viruses concerned have been confirmed and classified as zoonotic. Arenavirus haemorrhagic fever viruses can cause epidemics in some areas of the world. To assess the potential of similar viruses to become zoonotic in the future, bioinformatics of relatively conserved virus proteins, such as the L polymerase of arenaviruses, can be used to identify homology with viruses that might give future outbreaks. In this paper, new and archival metazoan transcriptome sequence data was used as a resource to identify similar sequences to known arenavirus sequences for the purpose of risk prediction. In essence, bioinformatics was utilized to provide a better understanding of the potential evolution and natural history of uncharacterized virus sequences in the GenBank database. Several matches were identified which, along with a reasoned approach to their phyla, was used to provide a likelihood score of their zoonotic potential.

## Introduction

There are many such viruses including arenavirus haemorrhagic fever viruses, SARS and MERS coronaviruses, Ebola virus, Marburg virus, Hendra virus and Nipah virus and the, all categorized as zoonotic RNA viruses, which can cause epidemics (Ehichioya et al., 2010; Fichet-Calvet and Rogers, 2009). *Arenaviridae* are a family of viruses composed of three genera, *Mammarenavirus, Reptarenavirus* and *Hartmanivirus* (Siddell et al., 2019). Mammarenaviruses distributing worldwide and cause febrile diseases for both humans and animals (Buchmeier et al., 2007). Reptarenaviruses are infected snakes (Hepojoki et al., 2015), were the infection in human have not been reported. Arenaviruses have a segmented ambisense single strand RNA genome, which is typically grouped as negative sense RNA viruses (Buchmeier et al., 2007). The large segment named as L and the short segment named as S (Bodewes et al., 2013). The L segment encodes the viral RNA dependent RNA polymerase (RdRp) or L polymerase and the small zinc finger motif protein (Z), whereas the S segment encodes the nucleoprotein (NP) and glycoprotein precursor (GPC) (Salvato et al., 1989).

The viral RNA dependent RNA polymerase (L polymerase) is the largest protein encoded by arenaviruses with a size of around 6.6 kb in a total genome of ∼11kb (Zapata and Salvato, 2013). The L polymerase contains an endonuclease domain that plays a significant role in mRNA cap-snatching during viral transcription (Morin et al., 2010). As L polymerase contains several essential catalytic domains it represents a protein which would be expected to be conserved, at least in part, by viruses using a similar mode of replication. Because the L polymerase is highly conserved it provides a useful source of amino acid sequences, for example from the catalytic site and from the endonuclease domain, for homology based searches of the unannotated databases to assess how widespread these conserved features are, and if they are found in virus sequences that may be of pandemic concern.

## Material and Methods

Bioinformatics software and tools were used in this research. Online tools such as translated basic local alignment search tool known as translated blast (tBLASTn) on National Center for Biotechnology Information (NCBI) available on (https://blast.ncbi.nlm.nih.gov/Blast.cgi) was used for searching translated nucleotide sequencing database that available in GenBank were Transcriptome ShoutGun Assembly (TSA) was used for reciprocal BLAST (McGinnis and Madden, 2004). Also, the open reading frames (ORFs) of TSA database were identified by the ExPASy translation tool (Gasteiger et al., 2003) on (http://web.expasy.org/translate/). Additionally, online tool kit including HHpred (Söding et al., 2005) that available online on (https://toolkit.tuebingen.mpg.de/#/tools/hhpred) were also utilized for protein homology detection and structure prediction of ORFs of TSA database. All identified proteins sequences were aligned by multiple sequence alignments (MSA) using Clustal Omega version (1.2.4) (McWilliam et al., 2013) available on (http://www.ebi.ac.uk/Tools/msa/clustalo/). Finally, MrBayes interface version 3.2.6 (Ronquist et al., 2011) was used for designing phylogenetic tree for L protein sequencing data of arenaviruses, TSA database and some other viruses. FigTree v1.4.3 software on (http://tree.bio.ed.ac.uk/software/figtree/) was used for viewing final phylogenetic tree of protein sequences that utilized at this research.

## Results

### Translated blast (tBLASTn) of arenaviruses

In this study, tBLASTn (McGinnis and Madden, 2004) facility on National Center for Biotechnology Information (NCBI) was used to align a query protein sequence to a nucleotide database assuming that any alignment could be in any possible reading frames translatable from any nucleic acid sequence. As bait, the L polymerase sequences of two species of mammarenavirus, Tacaribe virus (TCRV) and lymphocytic choriomeningitis virus (LCMV), as well as two species of reptarenavirus, Golden Gate virus (GGV) and California Academy of Science virus (CASV) were used to find similarities in the Transcriptome Shotgun Assembly (TSA) database. At least six genes sequences of different species were found to match to some part of the L polymerase of arenavirus species indicating some similarity with the arenaviruses used as the probe (Table 1).

**Table 1:** Hits identified following tBLASTn of L polymerase of arenaviruses. L polymerase of TCRV: Tacaribe virus (GenBank accession number NP_694848.1), LCMV: lymphocytic choriomeningitis virus (NP_694845.1), GGV: Golden Gate virus (YP_006590089.1), CASV: California Academy of Science virus (YP_006590093.1) were used for searching translated nucleotide in GenBank database by using translated blast (tBLASTn) were Transcriptome Shotgun assembly (TSA) allocated at search set of tBLASTn. The table shows the percentage of protein identity, E-value and accession number.

| **Virus** | **Transcribed RNA of TSA alignment** | **Number of hits (E-value)** | **Identity** | **Accession of TSA** |
| --- | --- | --- | --- | --- |
| TCRV | *Channa punctata* RP_73146 | 9e-05 | 41% | GEKU01073124.1 |
|  | *Catostomus commersonii* Contig_131767 | 0.001 | 51% | GECX01131681.1 |
| LCMV | *Channa punctata* RP_73146 | 7e-06 | 43% | GEKU01073124.1 |
|  | *Asymmetron lucayanum* comp27398_c0_seq1 | 6e-05 | 26% | GETC01011646.1 |
|  | *Catostomus commersonii* Contig_163647 | 0.002 | 36% | GECX01163539.1 |
|  | *Rhizopus oryzae* KNUBEL_RO_010568 | 0.18 | 24% | GDUK01010546.1 |
| GGV | *Channa punctata* RP_73146 | 5e-15 | 44% | GEKU01073124.1 |
|  | *Asymmetron lucayanum* comp27398_c0_seq1 | 4e-04 | 24% | GETC01011646.1 |
|  | *Catostomus commersonii* Contig_131767 | 0.002 | 48% | GECX01131681.1 |
|  | *Rhopilema esculentum* c77127_g1_i1 | 0.068 | 23% | GEMS01099264.1 |
| CASV | *Channa punctata* RP_73146 | 8e-13 | 37% | GEKU01073124.1 |
|  | *Catostomus commersonii* Contig_131767 | 0.001 | 35% | GECX01131681.1 |
|  | *Talitrus saltator* comp102738_c0_seq1 | 0.005 | 27% | GDUJ01083694.1 |
|  | *Asymmetron lucayanum* comp27398_c0_seq1 | 0.11 | 24% | GETC01011646.1 |

The data shows there are transcribed RNA sequences from extant species that align with the bait L polymerase structures including from *Channa punctate* (spotted snakehead fish), *Catostomus commersonii* (white sucker fish), *Asymmetron lucayanum* (lancelet), *Rhizopus oryzae* (fungus), *Rhopilema esculentum* (flame jellyfish) and *Talitrus saltator* (sand hopper). All these candidates had stretches of homology with similarity given an E-value <1. As the E-value indicates the number of the hits during database searching that are expected by chance the more significant E-values are less than 1 or zero whilst an E-value equal to 1 is not considered since it indicates only a chance alignment (Pearson, 1995; Pearson and Lipman, 1988). The target transcripts found (Table 1) were translated to allow further homology detection and structure prediction with other proteins found in the annotated protein database. Subsequently, the translated protein sequences from the TSA database and the sequences of mammarenavirus (TCRV and LCMV) and reptarenavirus (GGV and CASV) used as bait for arenavirus like isolates.

### Homology detection and structure prediction of identified proteins

The transcribed RNA of the TSA database (Table 1) were used to deduce protein homology with known proteins or domains using the SIB Bioinformatics Resource Portal (Gasteiger et al., 2003). The largest one or two segments of the open reading frame (ORFs) found were selected and used to find protein homology among other viruses in the database by using the online bioinformatics tool HHpred, version HHsuite-2.0.16mod (Söding et al., 2005). Interestingly, the HHpred output showed all the translated protein fragments of *Channa punctate, Catostomus commersonii, Asymmetron lucayanum, Rhizopus oryzae* and *Talitrus saltator*, identified originally as having homology to the arenavirus L polymerase, to have some homology with the La Crosse bunyavirus polymerase as found in structure 5amr-A in complex with the 3’ viral RNA. In some cases, homology was also apparent with Influenza C virus RNA-dependent RNA polymerase (5d98-B), whereas the *Rhopilema esculentum* sequence had homology with the structure of the L polymerase of vesicular stomatitis virus (5a22-A). The amount of protein homology varied among the translated proteins. For example, the protein homology of the *Rhizopus oryzae* sequence with La Crosse Bunyavirus polymerase (5amr-A) is the longest, reaching 854 amino acids, while the homology between the translated protein of *Catostomus commersonii* with Influenza C virus RNA-dependent RNA polymerase (5d98-B) was only 56 amino acids (Figure 1, 2, 3, and 4). Therefore, ORFs detected in the translated proteins of the transcribed RNA of TSA database showed homology with other viruses of known pathogenicity and could assist in the prediction the phylogenetic relationship among arenaviruses and other viruses archived in the GenBank databases.

**Figure 1:**
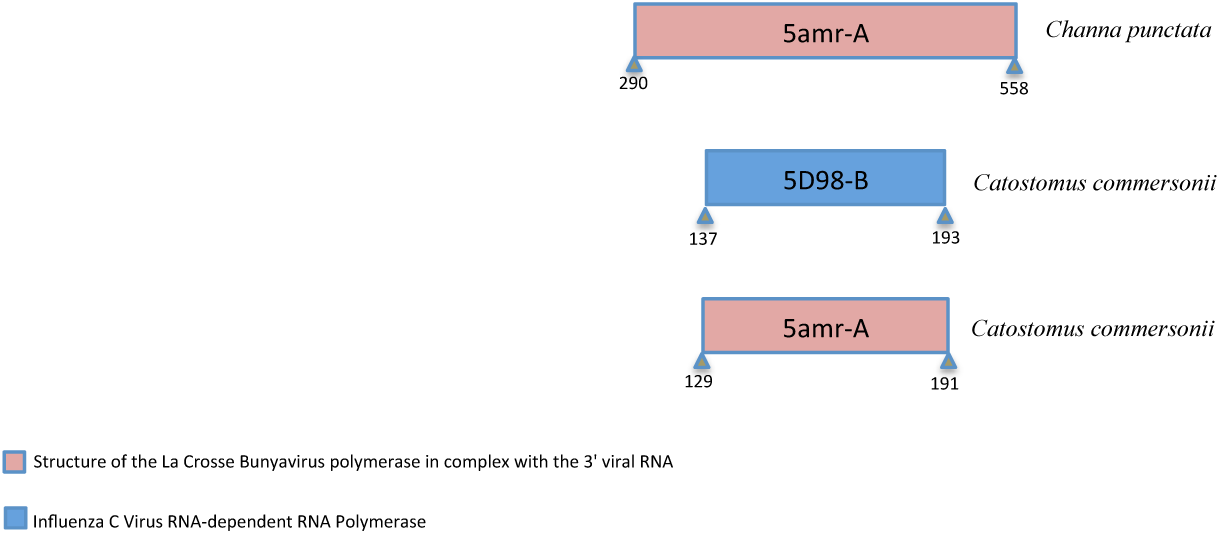
Homology detection of translated L polymerase of Tacaribe virus (TCRV). HHpred bioinformatics tools kit was used for protein analysis, while Microsoft PowerPoint was utilized for drawing the scales. The length of the scale depends on the gene size, the colour depends on virus gene homology. The 5amr-A (darksalmon colour) refers to structure of the La Crosse Bunyavirus polymerase in complex with the 3’ viral RNA, 5D98-B (blue colour) refers to Influenza C virus RNA-dependent RNA polymerase.

**Figure 2:**
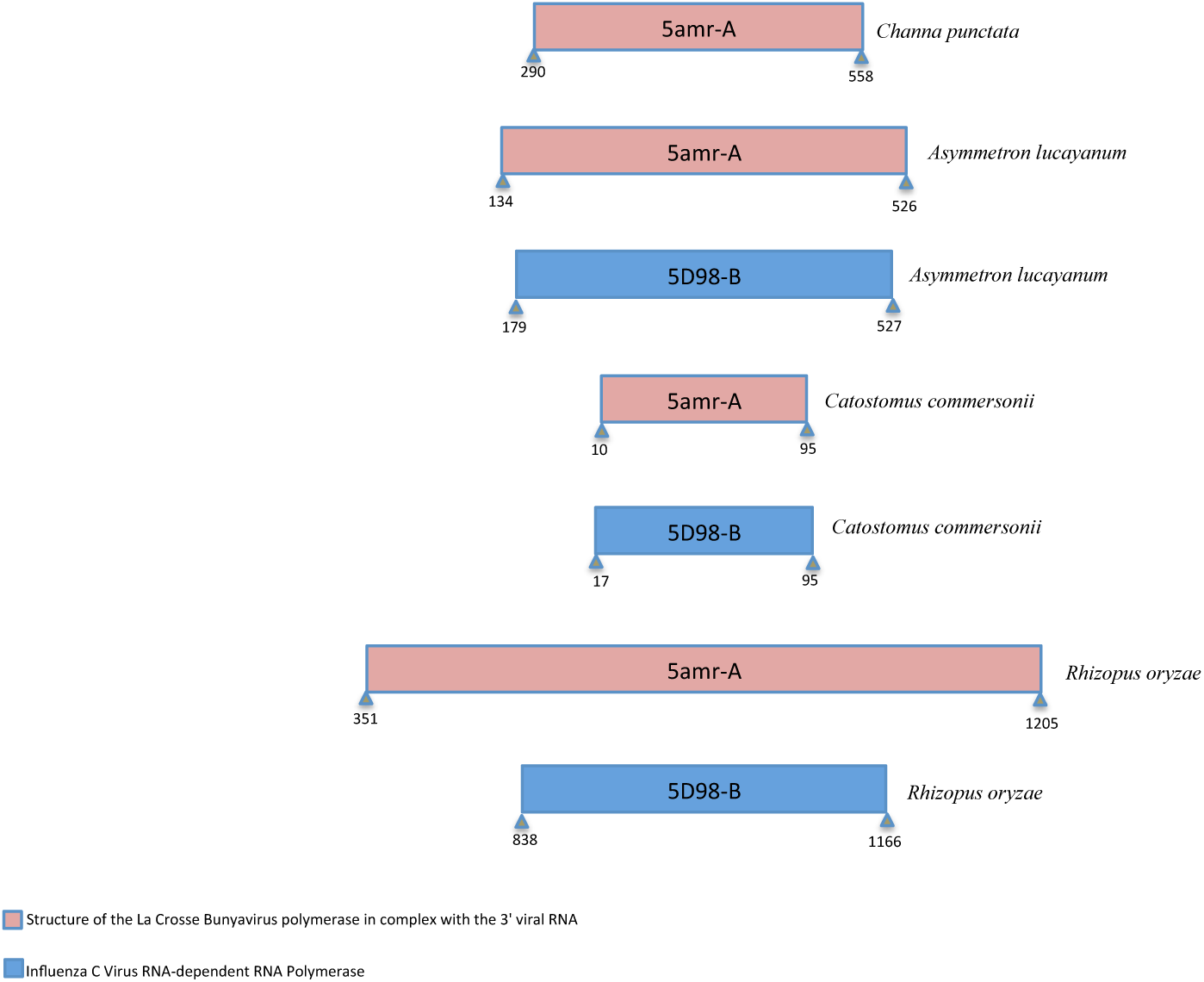
Homology detection of translated L polymerase of lymphocytic choriomeningitis virus (LCMV). HHpred bioinformatics tools kit was used for this purpose were Microsoft PowerPoint was used for drawing the scales. The length and the colour of scale depend on gene size and gene homology of virus. The 5amr-A (darksalmon colour): structure of the La Crosse Bunyavirus polymerase in complex with the 3’ viral RNA, 5D98-B (blue colour): Influenza C virus RNA-dependent RNA polymerase.

**Figure 3:**
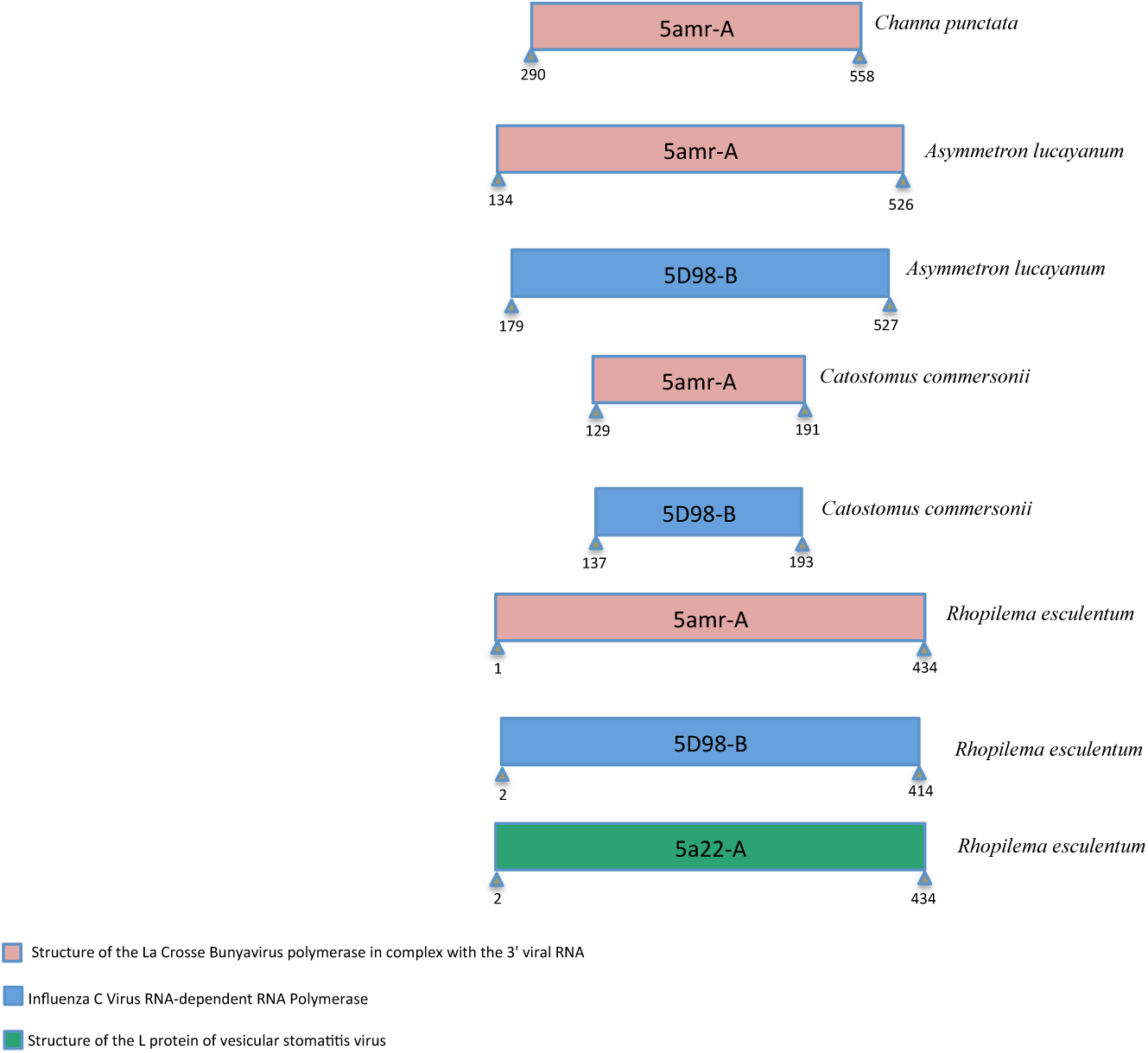
Homology prediction of translated L polymerase of Golden Gate virus (GGV). HHpred bioinformatics tools kit was used for protein analysis and Microsoft PowerPoint was used for drawing the scales. The scale in different colour and length depends on gene homology and gene size of the viruses, were 5amr-A (darksalmon colour) refers to structure of the La Crosse Bunyavirus polymerase in complex with the 3’ viral RNA, 5D98-B (blue colour) refers to Influenza C virus RNA-dependent RNA Polymerase, while the 5a22-A (green colour) refers to structure of the L polymerase of vesicular stomatitis virus.

**Figure 4:**
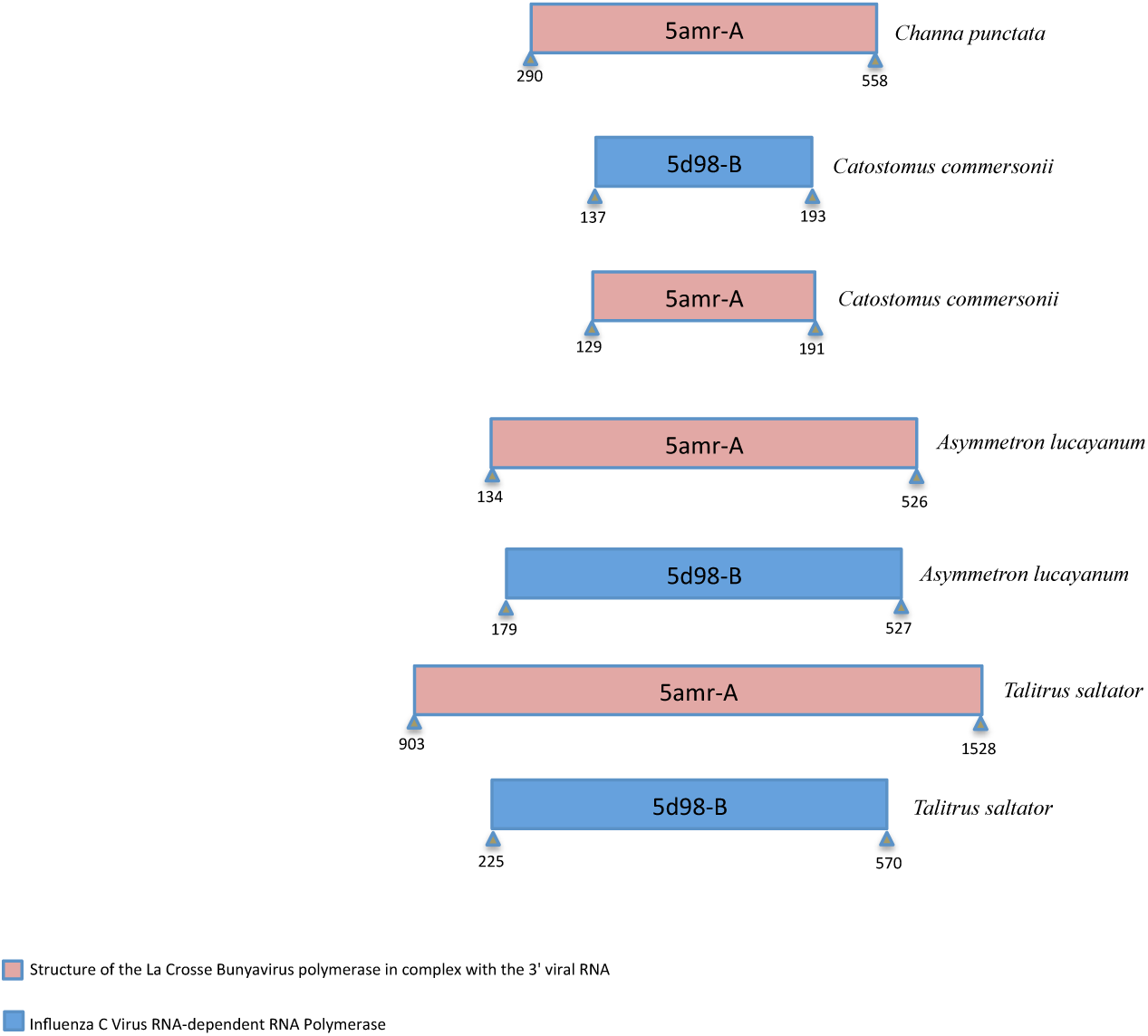
Homology detection and structure prediction of translated L polymerase of California Academy of Science virus (CASV). The translated protein was analysed by using HHpred bioinformatics tools, while the scales were drawn by using Microsoft PowerPoint. The scales showing in different colour and size, the colour refers to the identity of the gene, whereas the length refers to gene size of the target virus. The 5amr-A (darksalmon colour): structure of the La Crosse Bunyavirus polymerase in complex with the 3’ viral RNA, 5D98-B (blue colour): Influenza C virus RNA-dependent RNA polymerase.

### Multiple Sequence Alignment (MSA) of TSA database and arenavirus proteins

To evaluate the homology and conservation of the proteins found as well as study their phylogeny for the evolutionary relationships among the proteins of the arenaviruses and the TSA database, multiple sequence alignment (MSA) was used (Rani and Ramyachitra, 2016). The open reading frames (ORFs) sequence of the TSA database (Table 1) with the L polymerase of Tacaribe virus (TCRV), lymphocytic choriomeningitis virus (LCMV), Golden Gate virus (GGV) and California Academy of Science virus (CASV), were used for MSAs (Blazewicz et al., 2013). To do this, the sequence data of the translated proteins over the region of homology, as described in the results of the HHpred analysis, were aligned with L polymerases of (TCRV), (LCMV), (GGV) and (CASV) using Clustal Omega version (1.2.4). Partial MSAs as shown in Figure 5 were manually edited via Jalview 2.10.1 (Waterhouse et al., 2009).

**Figure 5:**
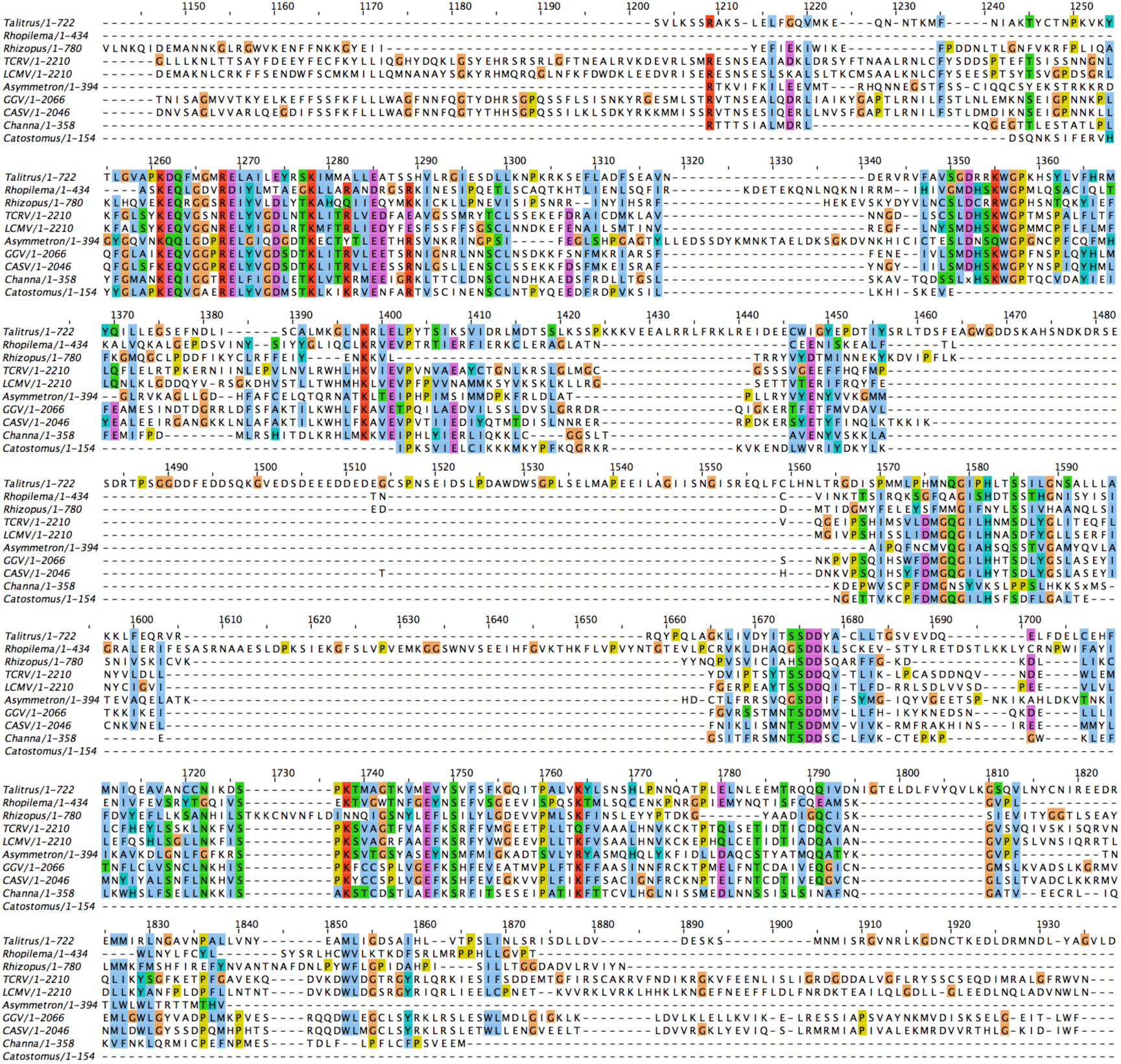
Multiple sequence alignment (MSA) of partial translated protein of TSA database hits with the L polymerase of arenaviruses. The translated proteins of *Channa punctate, Catostomus commersonii, Asymmetron lucayanum, Rhizopus oryzae, Rhopilema esculentum* and *Talitrus saltator* were aligned with L polymerase of arenaviruses including Tacaribe virus (TCRV), lymphocytic choriomeningitis virus (LCMV), Golden Gate virus (GGV) and California Academy of Science virus (CASV). The alignment starts around 1230 to 1870 amino acids, as the sequence of translated protein of TSA database matches with L polymerase of the arenaviruses species. Accession number of TRCV, LCMC, GGV and CASV were stated in Table 1. Clustal Omega (1.2.4) was used for the multiple alignments and Jalview 2.10.1(Waterhouse et al., 2009) was used for manual edition.

Figure 5 shows some shared regions of protein homology following multiple sequence alignments. In particular, two regions of L polymerase appeared to represent the most homologous stretches among the sequences aligned, residues ∼1210 to ∼1450 and residues ∼1560 to ∼1880. Interestingly the region 1050 – 1500 has been defined as the core RdRp in the Lassa L structure (Brunotte et al., 2011) while the later sequence also falls into a conserved region defined as domain IV based on arenavirus sequence analysis (Vieth et al., 2004). That these regions of conservation appear in such diverse targets suggests that it should be possible for design primers that could be used to amplify products from diverse sources as well as from current arenaviruses species by RT-PCR. In fact, RT-PCR was used for identify some species of mammarenavirus (Vieth et al., 2007) and reptarenavirus (Aqrawi et al., 2015). This could provide a diagnostic for the presence of the agents from which the TSA sequences were derived and also find other, related, virus sequences (Lozano et al., 1997; Vieth et al., 2005).

### Phylogeny analysis

To present a clear phylogenetic analysis between TSA database and arenaviruses that were used as probes, a phylogenetic tree was constructed that shows the relationship among these species. MrBayes version 3.2.6 (Ronquist et al., 2011) is a command line interface that was used for creating the trees by Bayesian Inference (BI). Bayesian Inference (BI) is based on the concept of probabilities that are based on prior expectation, after the analysis of some of the data. The L polymerase of arenaviruses sequences were aligned with the HHpred sequences from those TSA database hits found by using multiple sequence alignments (MSAs) by Clustal analysis (McWilliam et al., 2013), with some other viruses acting as roots for the phylogeny. The alignment file was then analysed by the MrBayes program. At a result, phylogenetic tree were visualized by a phylogenetic tree viewer program such as FigTree v1.4.3 software, (http://tree.bio.ed.ac.uk/software/figtree/) (Hall, 2018; Ronquist et al., 2011). In short, phylogenetic tree were successfully created by using Bayesian Inference (BI) facility and presented using FigTree v1.4.3 (Figure 6).

**Figure 6:**
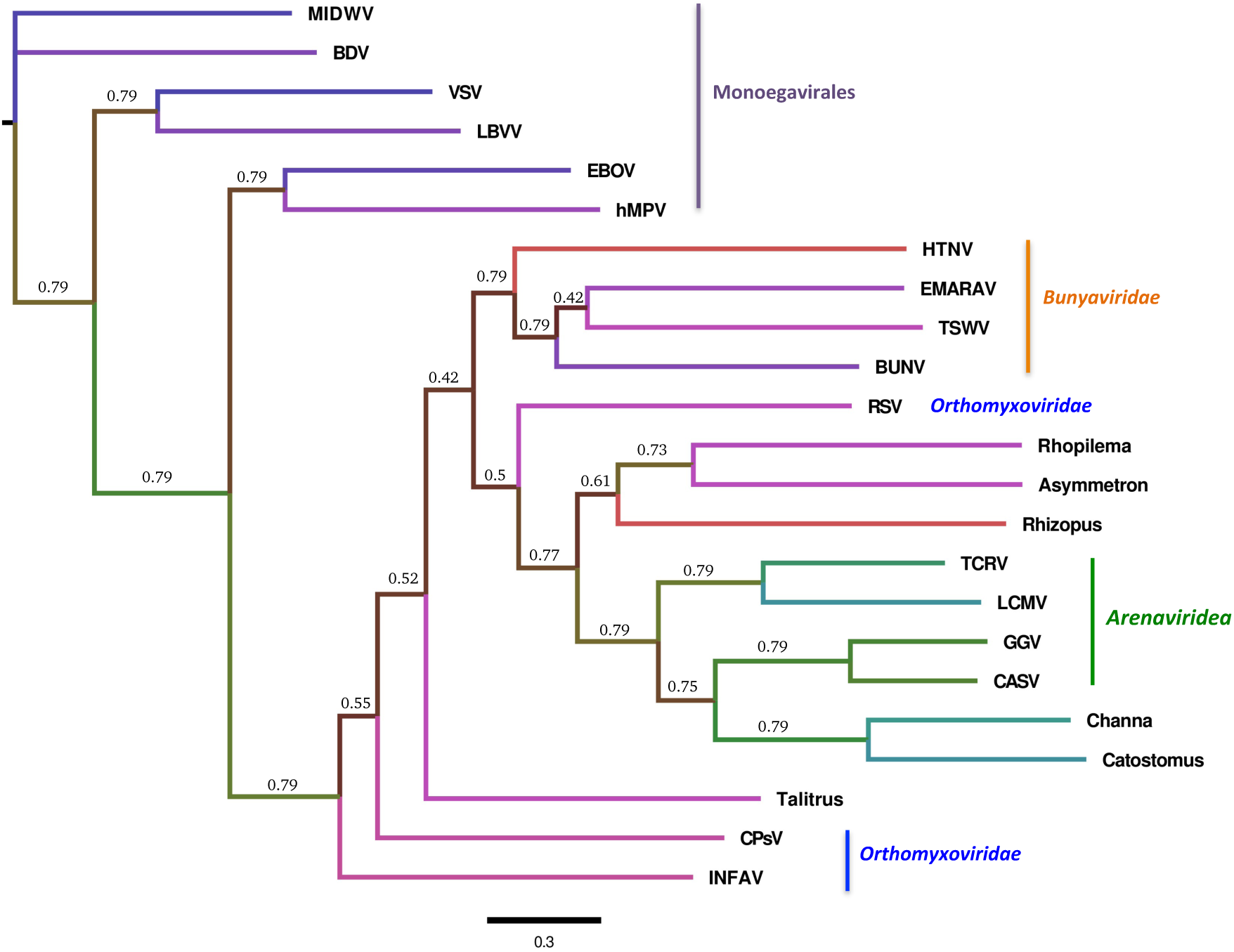
Molecular phylogenetic analysis by Bayesian Inference (BI). The phylogenetic tree was designed by using MrBayes v3.2.6 inference and manually edited by FigTree v 1.4.3. Average standard deviation of split frequencies was less than 0.01. The analysis involved 23 amino acid including translated protein sequences of TSA database, L polymerase sequences of arenaviruses and phylogeny root viruses protein sequences. Accession number of TRCV, LCMC, GGV and CASV were stated in Table 1. Accession number of root viruses, Hantaan virus (HTNV): NP_941982.1, Bunyamwera virus (BUNV): NP_047211.1, Rice stripe virus (RSV): AFM93820.1, Tomato spotted wilt virus (TSWV): AIA24440.1, Vesicular stomatitis virus (VSV): AAA48442.1, Citrus psorosis virus (CPsV): Q6DN67.1, European mountain ash ringspot-associated virus (EMARaV): Q6Q305.2, Lettuce big-vein associated virus (LBVaV): Q8B0U2.1, Influenza A virus: P03433.2, Midway nyavirus (MIDWV): YP_002905331.1, Borna disease Virus (BDV): CEK41892.1, Zaire ebolavirus (EBOV), AAG40171.1, Human metapneumovirus (hMPV): AII17600.1.

Viral RNA dependent RNA polymerase (vRdRp) is an ancient enzyme (Černý et al., 2014) which also, via its mutation rate, plays an important role in the evolutionary strategy of RNA viruses, leading to novel emerging viruses (de Farias et al., 2017; Vignuzzi et al., 2006). Viral genes and proteins evolve fast when compared to their cellular counterparts, a function of the vRdRp in driving their evolution (Cabanillas et al., 2013; Pickett et al., 2011; Smith et al., 2012) and Bayesian analysis of RNA viruses, including emerging viruses and others that were archived in GenBank database, is convenient for reconstruction the evolutionary distance between them. As demonstrated in Figure 6, the phylogram generated here suggests some evolution history between arenaviruses and the TSA database hits and with some other viruses that act as phylogeny tree roots including some viruses from *Orthomyxoviridae, Bunyaviridae* and *Monoegavirales*. To clarify, the arenaviruses species locate in the middle of the tree between *Rhizopus oryzae* and *Channa punctate, Asymmetron lucayanum* and *Rhopilema esculentum*. The bootstrap shows some differences in values between the viruses including arenaviruses and the species found in the TSA, while the values are the same between the other viruses. For instance, the bootstrap values between the arenavirus related TSA sequences from *Channa punctate* and *Catostomus commersonii* are the same and each around 0.79, which could indicate an evolutionary relationship between them.

## Discussion

The results of this study shown that when the viral RNA dependent RNA polymerase (L protein) of Tacaribe virus (TCRV), lymphocytic choriomeningitis virus (LCMV), Golden Gate virus (GGV) and California Academy of Science virus (CASV) were utilized for searching the translated nucleotides that are archived in GenBank in the Transcriptome Shotgun Assembly (TSA) database by tBLASTn, some distant homologues were found. The TSA database findings indicated sequences in six living species including *Channa punctate, Catostomus commersonii, Asymmetron lucayanum, Rhizopus oryzae, Rhopilema esculentum* and *Talitrus saltator* that have gene homology with the L polymerase of four species of arenavirus used for this study. When the translated proteins of these open reading frames (ORFs) from the TSA data were converted to amino acid sequences the translated proteins of the TSA database were found to have homology with the La Crosse Bunyavirus polymerase (5amr-A), Influenza C virus RNA-dependent RNA Polymerase (5d98-B) and with the L polymerase of vesicular stomatitis virus (5a22-A). Therefore, It was found that the L polymerase of some species of arenaviruses had significant TSA alignment with other species which indicates conserved genes among even distant arenaviruses.

Interestingly, phylogenetic tree alignment of all the protein sequences showed that the four arenavirus sequences located between *Rhopilema esculentum* at the distal end and *Talitrus saltator* at the bottom of the groups of the tree. The tree also shows that the translated protein of *Rhizopus oryzae* located between L polymerase of arenaviruses and the translated proteins of *Rhopilema esculentum* and *Asymmetron lucayanum*.

In summary, an unfiltered screen of the TSA database using conserved domains of known haemorrhagic mammarenaviruses and snake reptarenaviruses, revealed a number of homologies in different species in the database. It is not known if these sequences derive from viable replicating viruses but of the hits found several are unlikely as possible emerging viruses based on the hosts in which the sequences were found. Homologies found in *Rhizopus oryzae* (a fungus) are unlikely to be able to replicate in human cells despite this organism being an opportunistic pathogen. Similarly, sequences found in *Asymmetron lucayanum* (Amphioxides) are too distant to assume replication could occur in human cells. Initially, sequence homologies found in *Talitrus saltator* (a sand fly) might be considered as a possible arbovirus but this is unlikely as this is a non-human biting species so the opportunity for transmission is limited. However, protein sequence homologies found *Channa punctate* (fish), *Rhopilema esculentum* (jellyfish) and *Catostomus commersonii* (fish), could be plausible as components of viruses that at least had the opportunity to cause infection following human contact (by bite, sting and consumption respectively). These studies expand the range of possible viruses associated with new species and offer an indicator of their potential for human transmission. However, the study remains theoretical and experimental evidence for the viruses concerned and for pathogen transmission is yet to be confirmed.

More bioinformatics are recommended to find homology among arenaviruses and other sequences of living species that were archived in GenBank database, particularly from the rapidly growing transcriptome database, in order to monitor future outbreaks by viruses similar to arenaviruses. As a result of next generation sequencing the Transcriptome Shotgun Assembly (TSA) database is updated continuously, adding more species were reptarenavirus infection could be present. Any homology found can be used to design primers capable of confirming such targets in the lab by cDNA synthesis from the tissues used for the TSA database to help prove the viruses concerned are extant. Only by more RT-PCR on tissue samples, more bioinformatics and more collaboration will the future work of this study be possible.

